# A metagenomic DNA sequencing assay that is robust against environmental DNA contamination

**DOI:** 10.1101/2021.11.22.469599

**Authors:** Omary Mzava, Alexandre Pellan Cheng, Adrienne Chang, Sami Smalling, Liz-Audrey Djomnang Kounatse, Joan Lenz, Randy Longman, Amy Steadman, Mirella Salvatore, Manikkam Suthanthiran, John R. Lee, Christopher E. Mason, Darshana Dadhania, Iwijn De Vlaminck

**Author notes:** These authors contributed equally (Omary Mzava, Alexandre Pellan Cheng, Adrienne Chang).

## Abstract

Metagenomic DNA sequencing is a powerful tool to characterize microbial communities but is sensitive to environmental DNA contamination, in particular when applied to samples with low microbial biomass. Here, we present contamination-free metagenomic DNA sequencing (Coffee-seq), a metagenomic sequencing assay that is robust against environmental contamination. The core idea of Coffee-seq is to tag the DNA in the sample prior to DNA isolation and library preparation with a label that can be recorded by DNA sequencing. Any contaminating DNA that is introduced in the sample after tagging can then be bioinformatically identified and removed. We applied Coffee-seq to screen for infections from microorganisms with low burden in blood and urine, to identify COVID-19 co-infection, to characterize the urinary microbiome, and to identify microbial DNA signatures of inflammatory bowel disease in blood.

## INTRODUCTION

Metagenomic DNA sequencing is a routinely used tool to characterize the genetic makeup and species composition of microbial communities. In addition, metagenomic DNA sequencing of clinical isolates is increasingly used for unbiased detection of microbial infection. Nonetheless, sample contamination by environmental DNA plagues these assays. DNA contamination unavoidably occurs to a degree during the process of sample preparation for DNA sequencing and is particularly problematic for samples that have a low biomass of microbial DNA that can easily be overwhelmed by contaminating DNA^1–3^.

Multiple solutions have been proposed to overcome the impact of DNA contamination on low biomass metagenomic sequencing. DNA contamination can be avoided to an extent by processing samples in a clean room facility^4,5^. However, this approach does not avoid contaminant DNA present in reagents. Other approaches are based on batch-correction algorithms that identify microbial species detected in negative controls^5,6^. These methods however, tend to overcorrect, eliminate sample-intrinsic species that are also common DNA contaminants, and make the incorrect assumption that sample contamination is perfectly reproducible across all samples in a batch. Here, we describe <u>Co</u>ntamination-<u>F</u>r<u>ee</u> metagenomic <u>seq</u>uencing (Coffee-seq), a metagenomic sequencing method that is robust against DNA contamination. Coffee-seq tags sample-intrinsic, non-contaminant DNA, before DNA isolation with a chemical label that can be recorded via DNA sequencing. Contaminating DNA that is introduced in the sample after this initial tagging step can then be identified and eliminated. Several biochemistries can be envisioned for the initial DNA tagging step. Here, we implement deamination of unmethylated cytosines via bisulfite salt treatment of DNA. This chemistry does not require the use of enzymes or DNA oligos and can be applied directly to clinically relevant samples, such as blood and urine, as demonstrated in this work. We present an analysis of the technical performance of Coffee-seq and describe proof-of-principle applications of Coffee-seq to identify viral and bacterial COVID-19 co-infection from blood, to screen for urinary tract infection (UTI), to characterize the urinary microbiome, to screen for infections with low burden and prevalence in the blood of patients that presented with respiratory symptoms at outpatient clinics in Uganda, and to identify microbial DNA signatures in the blood of patients with inflammatory bowel disease (IBD).

### Coffee-seq working principle

For the practical implementation of Coffee-seq, we tag DNA by bisulfite salt-induced conversion of unmethylated cytosines to uracils (**Fig. 1A**). Uracils created by bisulfite treatment are converted to thymines in subsequent DNA synthesis steps that are part of the DNA sequencing library preparation. After DNA sequencing, contaminating DNA introduced after tagging can then be identified based on the lack of cytosine conversion. Bisulfite conversion does not require the use of commercial enzymes or oligos that are a frequent source of DNA contamination, and we found that it can be applied directly to the original sample, before DNA isolation. We developed a bioinformatics procedure to differentiate sample-intrinsic microbial DNA, contaminant microbial DNA, and host-specific DNA after Coffee-seq tagging (**Fig. 1B**, Methods). This procedure consists of three steps. First, host cfDNA is removed via mapping and k-mer matching. Given that CpG dinucleotides are heavily methylated in the human genome and rarely in microbial genomes, sequences containing CG dinucleotides are also removed. Second, remaining sequences that consist of more than three cytosines, or one cytosine-guanine dinucleotide are flagged and removed as likely contaminants. Last, a species-level filtering step is performed to remove any remaining reads that primarily originate from C-poor regions in the reference genome (**Fig. 1C**, Methods).

**Figure 1.**
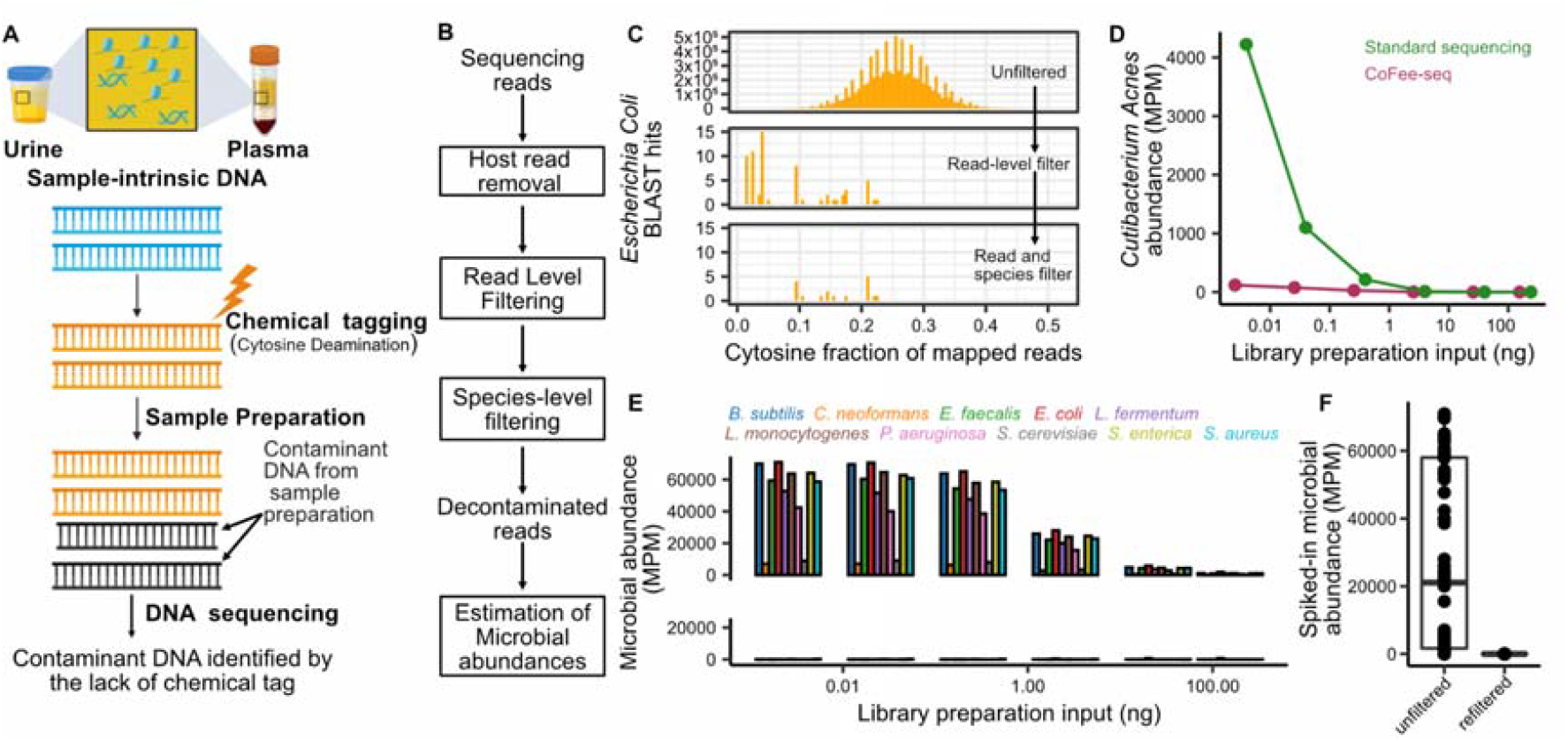
Coffee-seq proof-of-principle. **A)** Experiment workflow. Tagging of sample-intrinsic DNA by bisulfite DNA treatment is performed directly on urine or plasma. Contaminating DNA introduced after the tagging step is identified based on lack of cytosine conversion. **B)** Bioinformatics workflow. **C** Representative example of the cytosine fraction of mapped reads in an unfiltered (top) dataset, a read-level filtered dataset (middle) and a fully filtered dataset (bottom). **D)** Number of reads assigned to *Cutibacterium acnes* (common environmental DNA contaminant) inΦX174 DNA after conventional sequencing (green) and Coffee-seq (purple). **E)** Deliberate contamination assay. Detection of known contaminants before (top) and after (bottom) filtering. **F**) Number of reads assigned to contaminants.

We devised two assays to test the principle of Coffee-seq. First, we applied Coffee-seq and conventional DNA sequencing to samples of sheared ΦX174 DNA (New England Biolabs, #N3021S) with variable biomass (0.0025 ng, 0.025 ng, 0.25 ng, 2.5 ng, 26 ng, and 155 ng for Coffee-seq; 0.004 ng, 0.04 ng, 0.4 ng, 4 ng, 35 ng, and 240 ng for standard cfDNA sequencing). We first quantified the abundance of *Cutibacterium acnes (C. acnes)*, which is a frequent member of the normal skin flora and is routinely identified as a contaminant in DNA sequencing^7^. We observed an increase in *C. acnes* abundance with decreasing input biomass, as expected given that samples with a lower biomass are more susceptible to environmental contamination (**Fig. 1C**). We found that despite a ∼30% lower biomass at the beginning of library preparation for the Coffee-seq samples, far fewer *C. acnes* reads were present after Coffee-seq filtering (4223.8 and 119.5 MPM in the highest biomass samples, 1.48 and 0 MPM in the lowest biomass samples, before and after Coffee-seq filtering respectively; **Fig. 1D**).

Second, we performed Coffee-seq on sheared ΦX174 DNA samples with variable biomass (0.0025-155 ng; **Fig. 1E**) which we spiked after Coffee-seq tagging with 1 ng of sheared DNA from a well-characterized community of microbes to simulate microbial DNA contamination (10 species; Zymo Research, #D6305). Before applying the Coffee-seq bioinformatics filter, we observed a negative correlation between the ΦX174 DNA input biomass and the relative number of reads from the spike-in community, as expected (Pearson’s R = -0.54, p-value = 6.5×10^−6^; Spearman’s ρ = -0.82, p-value = 6.3×10^−16^; **Fig. 1E**). After applying the Coffee-seq filter, we observed an average percent decrease of 99.8% of molecules mapping to species of the spike-in community (**Fig. 1F**). Sequences mapping to *Escherichia coli* (*E. coli*) were the most abundant after filtering (58.89%). Given that ΦX174 genomic DNA is isolated after phage propagation in *E. coli* culture, we reasoned that these remaining reads were likely intrinsic to the original sample. Together, these experiments demonstrate the effectiveness of Coffee-seq for the detection and removal of DNA contaminants.

### Application of Coffee-seq to cell-free DNA in blood and urine

Cell-free DNA (cfDNA) in blood and urine has emerged as a useful analyte for the diagnosis of infection^8–15^. Metagenomic cfDNA sequencing can identify a broad range of potential pathogens with high sensitivity. Yet, because of the low biomass of microbial-derived cfDNA in blood and urine, metagenomic cfDNA sequencing is highly susceptible to environmental contamination, limiting the specificity of metagenomic cfDNA sequencing for pathogen identification.

To assess the performance of Coffee-seq in metagenomic cfDNA sequencing, we assayed a total of 169 cfDNA isolates (42 urine, 127 plasma) collected from five groups of subjects: **1)** 26 urine samples from a cohort of kidney transplant patients with and without UTI (16 UTI positive, 10 UTI negative; “kidney transplant cohort”), **2)** 16 urine samples collected early after transplantation from 10 kidney transplant patients that received a ureteral stent at the time of transplantation (samples were collected pre-stent and post-stent removal for 5 of the 10 patients; “early post-transplant cohort”), **3)** 56 plasma samples from a cohort of 44 patients presenting with respiratory symptoms at outpatient clinics in Uganda (28 sputum positive for Tuberculosis [TB], 16 sputum negative for TB; “Uganda cohort”), **4)** 41 plasma samples from a cohort of 32 patients diagnosed with IBD (16 patients with Crohn’s disease, 16 patients with ulcerative colitis; “IBD cohort”), and, **5)** 30 plasma samples from a cohort of 14 patients hospitalized with COVID-19 (“COVID-19 cohort”; see **Table S1** and Supplementary Information for details on the patients and samples included).

We performed Coffee-seq for all samples and obtained an average of 46.5 ± 23.6 million paired-end reads per sample. We detected and quantified the abundance of 68 genera that have been reported as frequent DNA contaminants in multiple independent studies (summarized in Ref. 4; **Fig 2A**, 49 of these genera detected in at least one sample). We found that 76% of these genera were completely removed from all samples after Coffee-seq filtering. We calculated the total number of molecules from all contaminant genera and observed an up to 3 orders of magnitude reduction after Coffee-seq filtering (reduced by a factor of 7.5, 1711.2, 177.6, 548.3, 547.2 for the kidney transplant cohort, early post-transplant cohort, Uganda cohort, IBD cohort, and COVID-19 cohort, respectively; **Fig. 2B**). We investigated the impact of Coffee-seq filtering on removing reads originating from the skin contaminant *C. acnes* (**Fig. 2C**). *C. acnes* was detected in all samples and completely removed from 50 samples by Coffee-seq filtering. In the remaining samples, we observed an up to 2 orders of magnitude reduction of *C. acnes* reads.

**Figure 2.**
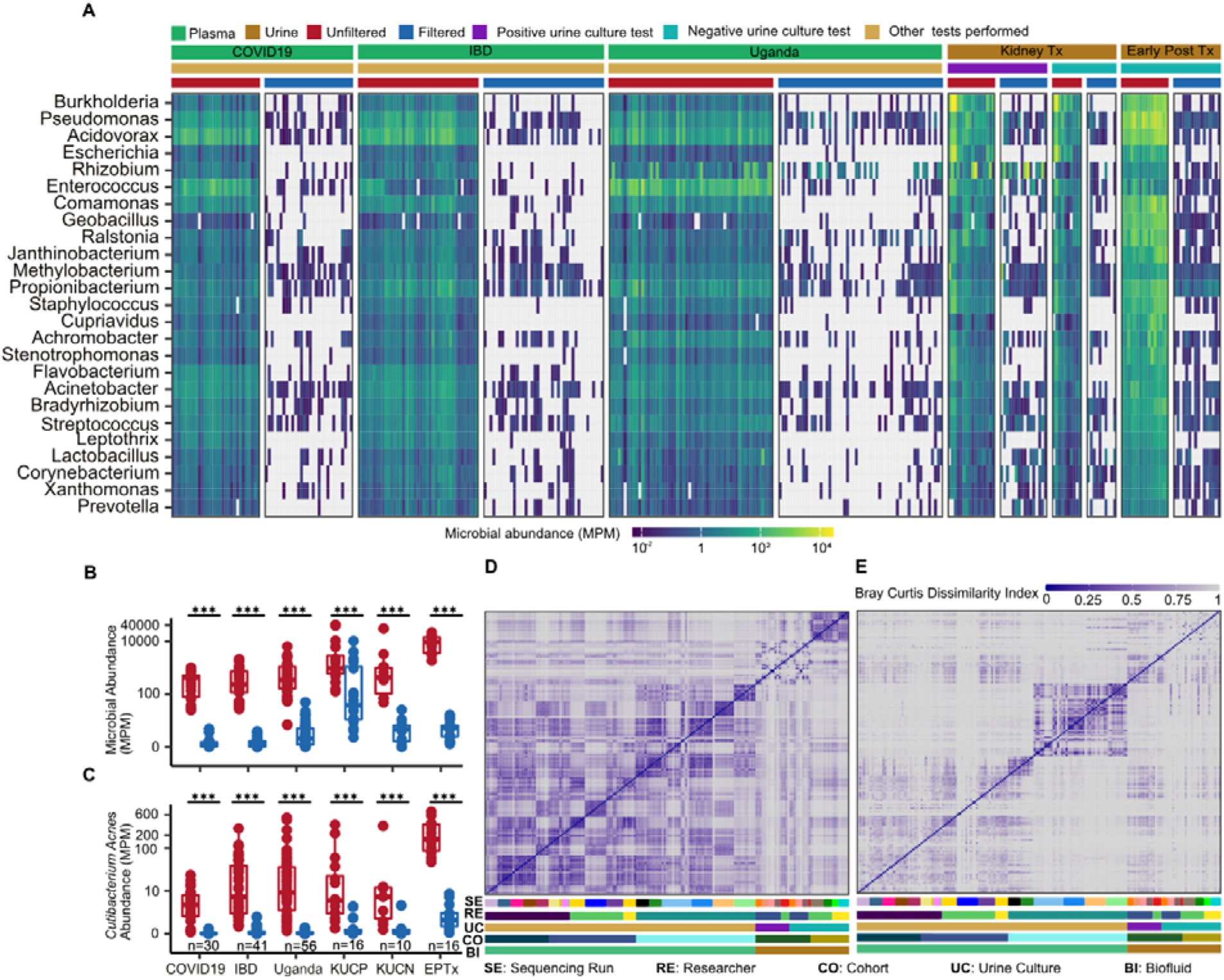
Coffee-Seq applied to cell-free DNA in urine and plasma. **A)** Microbial abundance of 25 most abundant common contaminant genera (selected from the 68 genera^4^) before and after Coffee-seq filtering in plasma and urine from five independent subject cohorts (Tx = transplant). Total abundance of all contaminant genera **B**) and *C. acnes* **C)** before and after Coffee-seq filtering (KUCP = Kidney Transplant cohort with positive urine culture, KUCN = Kidney Transplant cohort with negative urine culture, EPTx = Early Post Transplant cohort). Bray-Curtis dissimilarity index before **D**) and after **E**) filtering. Samples are organized by: sequencing batch, researcher performing the experiment, cohort, and biofluid. *** p-value < 0.001

We next evaluated the utility of Coffee-seq to correct for batch effects and to reveal true differences in microbiome profiles for different patient groups. To this end, we calculated the Bray-Curtis Dissimilarity Index for all clinical samples included in this study and sorted the datasets based on the following parameters: **1)** sequencing run, **2)** operator, **3)** urine culture test, **4)** study cohort, and **5)** biofluid type. Before Coffee-seq filtering, we observed a high similarity for samples assayed in the same experimental batches (**Fig. 2D**). Coffee-seq filtering removed these batch effects and revealed distinct cohort-specific microbiome profiles. Most notably, we observed distinct plasma microbiome profiles for plasma samples from the Uganda cohort (**Fig. 2E**). These results demonstrate that Coffee-seq directly applied to biofluids leads to a dramatic decrease in experimental noise and bias due to DNA contamination.

### Coffee-seq enables to screen for UTI and to characterize the urine microbiome

The healthy urinary tract was long believed to be sterile^16,17^, but this picture was challenged with recent advances in urine culture techniques that have identified bacteria in the urinary tract of both males and females^18^. Yet many microbes are difficult to cultivate *in vitro*, and bacterial culture can also be sensitive to contamination^19^. Therefore, comprehensive and accurate characterization of species colonizing the urinary microbiome is still lacking.

We reasoned that Coffee-seq could provide insight into the composition of the urine microbiome with both high sensitivity and specificity. We first applied Coffee-seq to 26 urine samples from 23 kidney transplant patients with and without infection of the urinary tract as determined by conventional urine culture (16 UTI positive [*Enterococcus faecalis*: n=3; *Enterococcus faecium*: n=1; *Escherichia coli*: n=10; *Klebsiella pneumoniae*: n=1; *Pseudomonas aeruginosa*: n=1] and 10 UTI negative). Coffee-seq consistently identified microbial cfDNA from species reported by urine culture (16/16 UTI positive samples; **Fig. 3A**). Coffee-seq also identified two Corynebacterium species (*Corynebacterium jeikeium* and *Corynebaterium urelyticum*) in one sample from a UTI positive patient (*E*.*coli*) with culture confirmed Corynebacterium co-infection. In addition, we found that samples from UTI positive patients had a significantly higher burden of total microbial DNA compared to samples from UTI negative patients (1451.8 ± 3024.7 MPM and 12.8 ± 17.6 MPM, respectively in the filtered samples; p-value = 1.1×10^−5^, Wilcoxon test; **Fig. 3B**). Conventional metagenomic sequencing (without Coffee-seq filtering) detected uropathogens with equal sensitivity but suffered from poor specificity: DNA from common uropathogens not identified by culture was detected in many samples, albeit with low abundance, including in samples from patients without UTI. We conclude that the improved specificity of Coffee-seq allows for more accurate characterization of co-infection networks in the scope of UTIs, and more accurate characterization of the normal urine microbiome in the absence of UTIs. It is important to note that two common skin microbes, *C. acnes* and *Staphylococcus epidermis*, were found in most samples (23/26 samples). While these two species have been shown to cause UTIs^20,21^, they may also have been introduced as contaminants at the time of urine collection, which underscores an important limitation of Coffee-seq: Coffee-seq is not robust against contamination that occurs before the tagging step.

**Figure 3.**
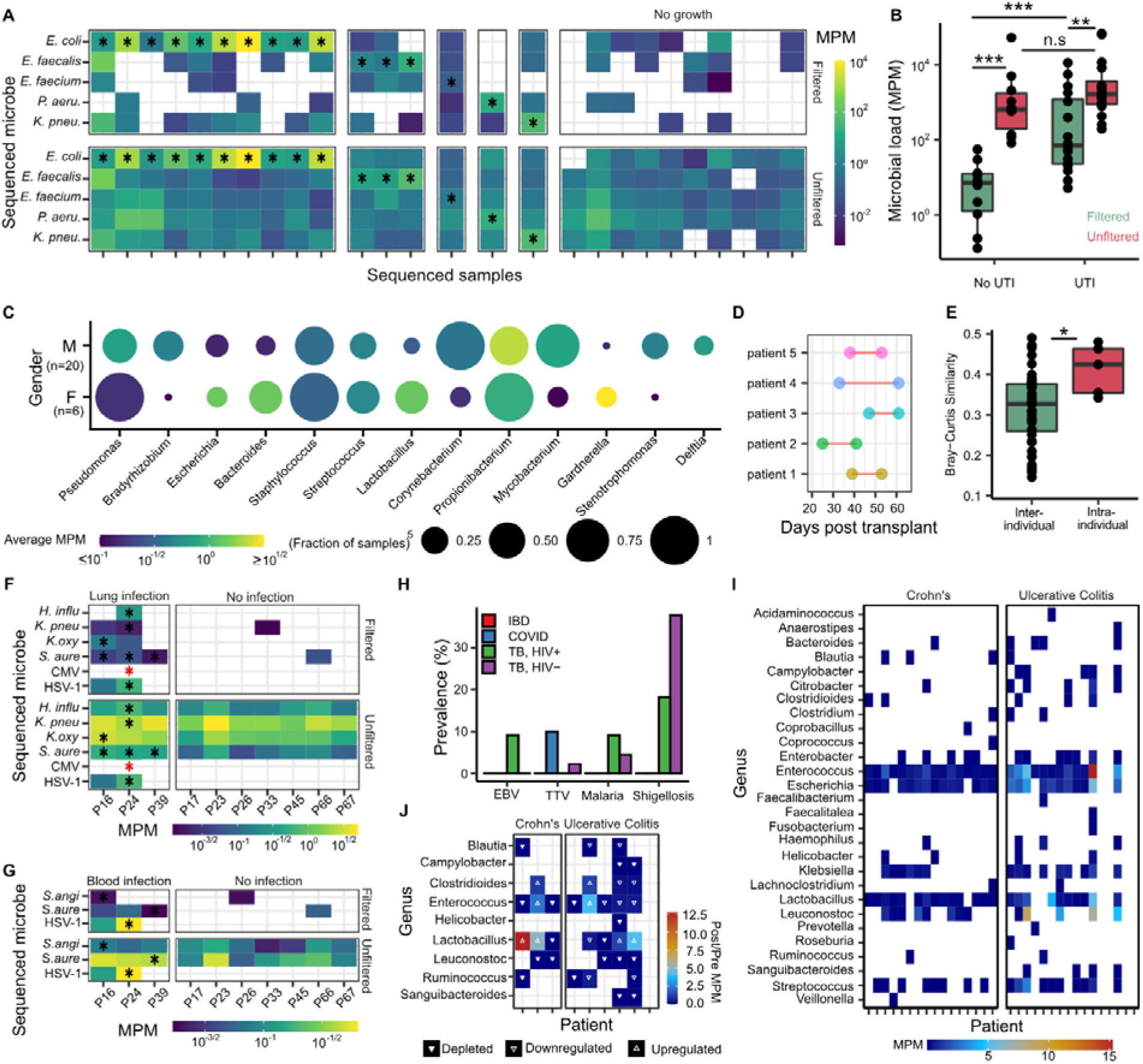
Application of Coffee-seq to plasma and urine. **A)** Heatmap of abundance of species (molecules per million, MPM) identified in patients with and without UTI, before and after application of Coffee-seq filter. **B)** Boxplot of the relative number of microbe-derived molecules (MPM) in samples from patients with and without UTI, before and after Coffee-seq filtering. **C)** Dot plot of the most abundant genera in urine from male and female kidney transplant recipients. **D-E)** Boxplot showing Bray-Curtis similarity index (as defined in **D**) of the urine microbiome within individual patients and between patients before and after stent removal. **F-G)** Heatmaps of the abundance of species identified in plasma from COVID-19 patients with and without culture confirmed **F)** lung and **G)** blood infection, before and after application of Coffee-seq filter (red * indicates detection by sputum culture only). Red boxes indicate positive culture tests. **H)** Barplot of the prevalence of Epstein-Barr Virus (EBV), Torque teno virus (TV), Malaria, or Shigellosis pathogens in different patient cohorts. **I)** Heatmap of the abundance of species identified in matched stool and plasma cfDNA samples in patients diagnosed with Crohn’s disease or ulcerative colitis. **J)** Heatmap of the change in abundance of gut specific bacteria before and after treatment. (Black * in panels A, F, and G indicates agreement with urine, respiratory and blood culture, respectively).

To explore the effect of gender on the urine microbiome, we analyzed isolates from culture confirmed UTI negative patients (n=26) from the kidney transplant (n=10) and early post-transplant (n=16) study cohorts (5 female, 14 male). This analysis yielded a small, but statistically insignificant, difference in total microbial load for male versus female patients (**Fig. S1**). We also observed that a subset of the most abundant genera was found in both male and female samples, with a marked variation in number of samples and abundances (**Fig. 3C**).

Studies investigating the temporal dynamics of urine microbiome in individuals can benefit from the high sensitivity and specificity achieved with our assay. We applied Coffee-seq to paired urine samples obtained from 5 kidney transplant patients collected at two time points before and after ureteral stent removal **(Fig. 3D)**. We compared the similarity of microbial composition between samples from the same patient (intra-individual) and between different patients (inter-individual) at different sampling points and observed that the microbial composition remained more similar in the same patient (**Fig. 3E)** than between different patients, supporting the utility of Coffee-seq to measure subtle dynamics in urine microbiome composition (Mean Bray-Curtis Similarity: 0.41±0.06 and 0.317±0.09 respectively, p-value = 3.1×10^−2^, Wilcoxon test).

### Coffee-seq identifies bacterial and viral co-infection of COVID-19 from blood

The COVID-19 pandemic is an unprecedented human health crisis. Viral or bacterial co-infection occurs in roughly 4% of hospitalized COVID-19 patients but can occur in up to 30% of COVID-19 patients admitted to the intensive care unit^22^. Co-infection has been associated with longer fever duration, and increased admittance to the intensive care unit and ventilation treatment^23^. We reasoned that Coffee-seq may offer sensitive detection of bacterial co-infection in COVID-19 patients with improved specificity over conventional metagenomic sequencing assays.

We applied Coffee-seq to 30 plasma samples from 14 patients with COVID-19 collected as part of a clinical study aimed at identifying predictors of disease severity. Respiratory and blood cultures were obtained as part of standard clinical care. Three patients (P16, P24, P39) tested positive for blood borne infection and respiratory tract infection, while all other patients were not diagnosed with COVID-19 co-infection. Coffee-seq identified the causative pathogen in 3/3 blood infection cases and 7/8 respiratory infection cases (**Fig. 3F-G**). Conventional metagenomic sequencing (without Coffee-seq filtering) was equally sensitive to these pathogens but was limited by specificity (**Fig. 3F-G**). Of interest, while we did not obtain plasma collected the day of infection for P24, we identified cfDNA originating from *K. pneumoniae* and *Haemophilus influenzae*, for which the patient tested positive four days later. While further investigation is necessary to resolve discrepancies between positive culture results and microbial cfDNA detection, these results suggest that Coffee-seq may be able to identify cases of infection earlier than traditional culture methods, and with improved specificity compared to conventional metagenomic sequencing techniques.

### Coffee-seq identifies bacterial and viral infections with low prevalence and low microbial burden

Neglected tropical diseases significantly impact the public health and economies of low-income countries. Treatments exist for many of these diseases, but development and deployment of reliable diagnostic tests has been slow^24^. We reasoned that Coffee-seq could be used to screen for infections with low prevalence and low microbial burden.

We applied Coffee-seq to 56 plasma samples from 44 individuals who presented with symptoms of respiratory illness at outpatient clinics in Uganda (28 sputum positive tuberculosis, 16 sputum negative tuberculosis). Nine of these individuals were HIV positive at the time of sample collection. We mined the data to determine the prevalence of infections endemic to Uganda and compared with results obtained for plasma samples collected from subjects that live in North America (54 plasma samples from the IBD cohort; 30 plasma samples from the COVID-19 cohort). We screened the samples for Epstein-Barr virus, Torque Teno virus, and pathogens associated with malaria (*Plasmodium vivax* and *P. falciparum*), and shigellosis (*Shigella sonnei, S. dysenteriae, S. boydii*, and *S. flexneri*). These pathogens were found at varying rates in samples from the Uganda cohort **(Fig. 3H)**: malaria (3/44), Epstein-Barr virus (1/44), shigellosis (19/44), and torque teno virus (1/44), but not in the IBD cohort. Torque teno virus, which has previously been reported to be elevated in immunocompromised patients^8^, was identified in 3/30 COVID-19 patient samples, all from patients who had received a bone marrow transplant prior to sample acquisition.

### Coffee-seq identifies signatures of bacterial translocation from the gastrointestinal tract

Bacterial translocation of intestinal microbes through mucosal membranes is believed to be a normal phenomenon, but has been found to occur more frequently in patients experiencing gut flora disruption^25,26^. In patients with inflammatory bowel disease, gut vascular barrier disruption has been linked to increased intestinal permeability and subsequent microbial translocation across the mucosal membrane^27,28^. The translocation of gut bacteria and their products to extraintestinal sites can result in systemic inflammation, resulting in autoimmune or other non-infectious diseases. Detecting signatures of translocation is therefore important but difficult in view of the low abundance of microbial DNA due to translocation in blood.

To identify signatures of bacterial translocation, we compared whole genome shotgun sequencing of fecal samples from 32 patients (Crohn’s n=16, ulcerative colitis, n=16) to matched plasma cfDNA samples assayed using Coffee-seq. We first quantified bacterial species identified in matched fecal and plasma samples (**Fig. 3I**). We identified cfDNA derived from gut-specific microbes in all patient samples, though to a much greater extent in individuals with ulcerative colitis (1.40±1.4 vs 6.82±10.6 MPM of gut specific bacteria for Crohn’s disease and ulcerative colitis, respectively). To investigate the effects of treatment on bacterial translocation, we collected additional stool and plasma samples from nine patients (Crohn’s n=3, ulcerative colitis n=6) after treatment initiation and performed whole genome shotgun sequencing of stool and Coffee-seq on plasma cfDNA. We quantified the relative abundance of gut-specific bacterial species before and after treatment and found that the burden of cfDNA decreased for most bacterial species (28/36) following treatment, which may be explained by a reduction in the degree of bacterial translocation with treatment (**Fig. 3J**). Of interest, out of seven subjects for which we detected *Lactobacillus* before treatment, five displayed an increase in *Lactobacillus* species burden in blood after treatment (up to 12.7-fold increase after treatment and an average of 3.36-fold MPM increase after treatment across all samples). *Lactobacillus* has been shown to promote gastrointestinal barrier function, protecting the gut from pathogenic bacteria and preventing inflammation^28^. For bacterial species besides *Lactobacillus*, we find an average of 0.3-fold MPM reduction after treatment. These preliminary results support the use of Coffee-seq to identify subtle signatures of bacterial translocation in the blood.

## DISCUSSION

We report Coffee-seq, a method for metagenomic DNA sequencing that is robust against DNA contamination. In contrast to prior methods for the management of DNA contamination that have relied on algorithmic batch correction or the use of known-template or no-template controls, Coffee-Seq uses a physical labeling technique to differentiate sample-intrinsic DNA from contaminating DNA. The principle of Coffee-seq has the potential for broad application in contexts where metagenomic analyses of isolates with low biomass of microbial DNA are required. In this proof-of-principle study, we have explored applications of Coffee-seq to quantify microbial cell-free DNA in human biofluids. Metagenomic sequencing of microbial cell-free DNA in blood or urine is a highly sensitive approach to screen for a broad range of viral or bacterial pathogens, but because of the low biomass of microbial DNA in blood and urine this method is highly susceptible to DNA contamination leading to a high false positive rate. We implemented Coffee-seq tagging of cell-free DNA in plasma and urine by bisulfite-induced deamination of unmethylated cytosines and show that this approach reduces background signals from common contaminants by up to three orders of magnitude. Coffee-seq thereby dramatically improves the specificity of metagenomic cfDNA analyses, opening up a broad range of applications, e.g. infectious disease with low microbial burden or syndromes that are accompanied by subtle changes in the plasma or urine microbiome.

In its current implementation, Coffee-seq has several limitations. First, Coffee-seq is only robust against DNA contamination introduced after the labeling step. We implemented Coffee-seq tagging directly on biofluids, which allowed us to identify contaminants introduced during DNA isolation or library preparation but not during the sample collection or isolation of the plasma from whole blood. Second, the specific labeling strategy we have implemented here inherently modifies the DNA sequence and thereby limits the resolution of sequence-based analyses. Alternative implementations of contamination-free sequencing that do not introduce sequence alterations can be considered. Last, the principles introduced here can be adopted for molecular assays beyond whole genome sequencing, including amplicon sequencing assays, e.g. 16S rRNA profiling, or PCR assays.

## METHODS

### Study Cohort and sample collection

#### Uganda cohort and sample collection

Forty-four plasma samples were collected from individuals seeking tuberculosis treatment in Uganda. Briefly, peripheral blood was collected in Streck Cell-Free BCT (Streck #230257) and centrifuged at 1600 x g for 10 minutes. Plasma was stored in 1 mL aliquots at -80°C. The study was approved by the Makerere School of Medicine Research and Ethics Committee (protocol 2017-020). All patients provided written informed consent.

#### IBD cohort sample collection

Peripheral blood samples were collected under IRB approved protocol (1806019340) at the Jill Roberts Center for IBD at Weill Cornell Medicine. PBMCs and plasma were fractionated using a Ficoll-Hypaque gradient.

#### Stool sample collection

DNA from fecal samples was isolated using the MagAttract PowerMicrobiome DNA/RNA kit with glass beads (Qiagen, Germany). Metagenomic libraries were prepared using the NEBNext Ultra II for DNA Library Prep kit (New England Biolabs, Ipswich, MA) following the manufacturer’s protocol. The DNA library was sequenced on an Illumina HiSeq instrument using a 2×150 paired-end configuration in a high output run mode.

#### COVID-19 cohort sample collection

Samples were collected as part of an observational study among individuals with COVID-19^29,30^ that were treated at New York Presbyterian Hospital and Lower Manhattan Hospitals, Weill Cornell Medicine. The study was approved by the Institutional Review Board of Weill Cornell Medicine (IRB 20-03021645), and informed consent was obtained from all participants.

#### UTI cohort sample collection

Twenty six urine samples were collected from 23 kidney transplant recipients who received care at New York Presbyterian Hospital–Weill Cornell Medical Center. The study was approved by the Weill Cornell Medicine Institutional Review Board (protocols 1207012730). All patients provided written informed consent. Patients provided urine specimens using a clean-catch midstream collection protocol. The urine specimen was centrifuged at 3000 *x g* for 30 minutes and supernatant was stored as 1 mL of 4 mL aliquots.

#### Early post transplant sample collection

Urine specimens collected within 10 ± 5 days of ureteral stent removal from patients who agreed to participate in the WCM IRB approved protocol # 20-01021269 were included in this study. Urine specimens were collected within 47 ± 11 days post-kidney transplantation. The presence of UTI was excluded by a negative urine culture and the absence of pyuria. This study was approved by the Weill Cornell Medicine Institutional Review Board (protocol 20-01021269).

#### Definition of Positive and Negative urine culture for the UTI and Early post-transplant cohorts

A positive urine culture was defined as a culture growing an organism identified to at least the genus level (≥10,000 cfu/mL). A urine culture was defined as negative when either no organism was isolated in culture (<1000□cfu/mL) or the organism was unidentified to either the genus or species level (i.e., unidentified) and the colony count was <10,000□cfu/mL.

##### Coffee-seq in plasma

An aliquot of 520 µL of plasma was centrifuged at 14,000 RPM for 10 minutes at 12°C to pellet cellular debris. The supernatant was transferred to a new 1.5 mL tube and the final volume was brought up to 1000 µL with PBS. The solution was heated to 98°C for 10 minutes and mixed at 1000 RPM to coagulate the albumin present in plasma. The solution was then centrifuged at 4000 RPM for 10 minutes. 500 µL of supernatant was transferred to 15 mL falcon tube containing 3.25 mL of ammonium bisulfite solution (Zymo Research, product #5030) and shaken in a thermomixer at 98°C for 10 minutes (15s on/30s off). Samples were then transferred to a thermomixer at 54°C for 60 minutes (15s on/30s off). Then, cfDNA extraction was performed using the QIAamp Circulating Nucleic Acid Kit using the 4-mL plasma protocol (Qiagen, product #55114). Prior to DNA elution, 200 µL of L-Desulphonation buffer (Zymo Research, product #5030) was added to the columns for 15 minutes, followed by two washes with 200 µL absolute ethanol. DNA was then eluted according to manufacturer recommendations, and single-stranded library preparation is performed (Claret Biosciences, product #CBS-K150B). Libraries were then sequenced on an Illumina sequencer.

##### Coffee-seq in urine

An aliquot of 520 µL of urine was centrifuged at 14,000 RPM for 5 minutes to pellet cellular debris. 500 µL of supernatant was transferred to a new 15 mL falcon tube containing 3.25 mL of ammonium bisulfite solution (Zymo Research, product #5030) and heated to 98°C for 10 minutes. Samples were then kept at 54°C for 60 minutes. Then, cfDNA extraction was performed using a commercially available column-based kit (Norgen Biotek, product #56700). Prior to DNA elution, 200 µL of L-Desulphonation buffer (Zymo Research, product #5030) was added to the columns for 20 minutes, followed by two washes with 200 µL absolute ethanol. DNA was then eluted according to manufacturer recommendations, and single-stranded library preparation was performed (Claret Biosciences, product #CBS-K150B). Libraries were then sequenced on an Illumina sequencer.

##### Alignment to the human genome

Adapter and low quality bases from the reads were trimmed using BBDuk^31^ and aligned to the C-to-T and G-to-A converted human genome using Bismark^32^ (Bismark-0.22.1). PCR duplicates were removed using Bismark.

##### Depth of coverage

The depth of sequencing was measured by summing the depth of coverage for each mapped base pair on the human genome after duplicate removal, and dividing by the total length of the human genome (hg19, without unknown bases).

##### Bisulfite conversion efficiency

We estimated bisulfite conversion efficiency by quantifying the rate of C[A/T/C] methylation in human-aligned reads (using MethPipe^33^ V3.4.3), which are rarely methylated in mammalian genomes.

##### Metagenomic abundance estimation from sequencing data

Metagenomic analysis is performed as previously described^12^. Specific to Coffee-seq, read-level filtering of contaminants is performed by removing sequenced reads with 4 or more cytosines present, or one methylated CpG dinucleotide (the latter represents unmapped, human-derived molecules). Species-level filtering based on the distribution of mapped reads is carried out by first aligning filtered and unfiltered datasets independently. Cytosine-densities of mapping-coordinates in both datasets are measured using custom scripts, and their distributions are compared using a Kolmogorov-Smirnov test. Significantly different filtered-unfiltered distributions are further processed (D-statistic > 0.1 and p-value < 0.01). Briefly, filtered datasets whose distribution of cytosines at mapped locations is significantly lower than unfiltered datasets have one read removed, and are re-tested for differences in their distribution. If the distributions are more similar (as measured through the same criteria), it is filtered out. This process is repeated until distributions are no longer significantly different, or if all reads are removed. Metagenomic abundances of filtered datasets are estimated using GRAMMy as previously described in Ref 12. Microbial abundance in downstream analyses was quantified as Molecules Per Million reads (MPM).

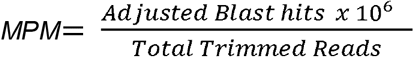

### Identification of translocated gut bacteria in plasma

Fecal shotgun metagenomic data for 41 samples was obtained from 32 patients diagnosed with inflammatory bowel disease (IBD). Low-quality bases and Nextera-specific sequences were trimmed (Trim Galore). Reads were aligned (Bowtie2^34^) against the human references (UCSC hg19). Unaligned reads were extracted and assembled with metaSPAdes^35^ and classified with Kaiju^36^.

Paired cfDNA samples were filtered as previously described and aligned to the assembled reads with Bismark. Mapped reads with a minimum quality score of 15 were extracted and filtered for gut-specific microorganisms identified by The Human Gut Microbiome Atlas^37^.

### Statistical analysis

All statistical methods were performed in R version 4.0.5. Groups were compared using a two-sided Wilcoxon Rank Sum test. Boxes in the boxplots indicates 25th and 75th percentile, the band in the box indicated the median and whiskers extend to 1.5 x Interquartile Range (IQR) of the hinge.

### Code and Data Availability

All scripts used in this study are available at https://github.com/omrmzv/CoffeeSeq.ΦX174 DNA sequencing data used in the proof of principle experiments has been deposited in NCBI’s Sequence Read Archive (SRA) under Bioproject ID (PRJNA782310). Sequencing data from human plasma cfDNA will be deposited in the database of Genotypes and Phenotypes (dbGaP)

## ACKNOWLEDGMENTS

We thank the Cornell Genomics Center for help with sequencing assays, the Cornell Bioinformatics facility for computational assistance and Michael Satlin and Lars Westblade for helpful discussions. A special thanks to Dr. Alfred Andama for his research supervision with the Infectious Disease Research Collaboration (IDRC) for samples collected and characterized in Kampala, Uganda. This work was supported by R01AI146165 (to I.D.V.), R21AI133331 (to I.D.V.), R21AI124237 (to I.D.V.), DP2AI138242 (to I.D.V.), R01AI151059 (to I.D.V., J.R.L., M.S., D.D.), R37 AI051652 (to M.S), a Synergy award from the Rainin Foundation (to I.D.V. and R.L.), a grant from the Bill and Melinda Gates Foundation INV-003145 (to I.D.V.). A.C. was supported by the National Institutes of Health under the Ruth L. Kirschstein National Research Service Award (6T32GM008267) from the National Institute of General Medical Sciences. A.P.C. is supported by a National Sciences and Engineering Research Council of Canada PGS-D3 fellowship.

## CONFLICTS

IDV, OM, APC and AC have submitted a patent related to the present work. APC, IDV, DD, and JRL are inventors on the patent US-2020-0048713-A1 titled “Methods of Detecting Cell-Free DNA in Biological Samples.” I.D.V. is a member of the Scientific Advisory Board of Karius Inc., Kanvas Biosciences and GenDX. JRL received research support under an investigator-initiated research grant from BioFire Diagnostics, LLC.

