## Supplementary material for "A metagenomic DNA sequencing assay that is robust against environmental DNA contamination": Table S1

*These authors contributed equally.

**Table S1**. Overview of all the samples included in this study

| **Cohort** | **Biofluid** |  | **Patients** | **Samples** |
| --- | --- | --- | --- | --- |
| Kidney transplant | Urine |  | 23 | 26 |
|  |  | *UTI+* |  | *16* |
|  |  | *UTI-* |  | *10* |
| Early post-transplant | Urine |  | 10 | 16 |
|  |  | *Paired pre/post stent removal* | 5 | 10 |
| Uganda | Plasma |  | 44 | 56 |
|  |  | *Sputum positive* | 28 |  |
|  |  | *Sputum negative* | 16 |  |
|  |  | *HIV+* | 9 | 11 |
|  |  | *HIV-* | 35 | 45 |
| COVID-19 | Plasma |  | 14 | 30 |
| Inflammatory bowel disease | Plasma |  | 32 | 41 |
|  |  | *Ulcerative colitis* | 16 | 22 |
|  |  | *Crohn’s Disease* | 16 | 19 |
|  |  | *Paired pre/post therapy* | 9 | 18 |
|  |  | *Matched whole genome sequenced fecal samples* | 32 | 41 |

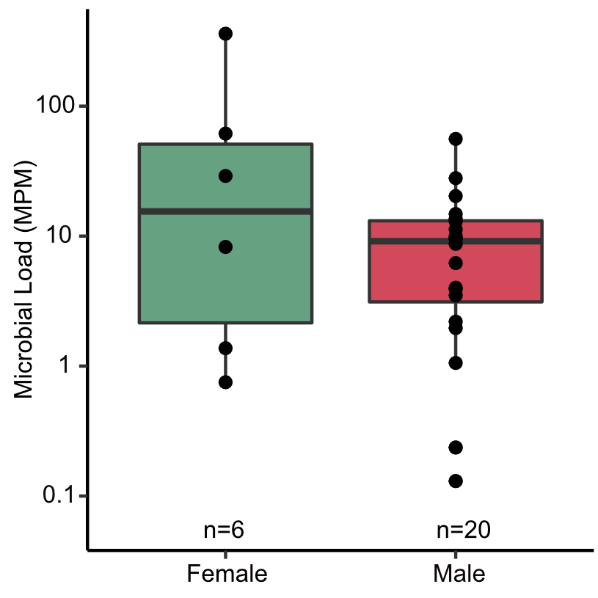

**Supplementary Figure S1**: Microbial load variation measured by Coffee-seq, as a function of gender

**Supplementary Methods**

### **Coffee-seq in Urine**

#### ***Reagents***

- Lightning Conversion Reagent (Zymo Cat# D5032-1)
- L-Desulphonation Buffer (Zymo Cat# D5030-5)
- Urine Cell-Free Circulating DNA Purification Midi Kit (Norgen Cat #56700)
- UDI Primer Set-24 (SRSLY Cat# CBS-UD-24)
- PicoPlus DNA NGS Library Preparation Base Kit (SRSLY Cat# CBS-K250B-24)
- AmpureXP Beads (Beckman Coulter Cat# A63880)
- KAPA HiFi HotStart Uracil+ ReadyMix (2X) (Roche Cat# 7959052001)
- Qubit ssDNA Assay Kit (Invitrogen Cat# Q10212)

#### ***Equipment***

- Two thermoshaker with 15 mL adapters
- Water bath
- Centrifuge

#### ***Step 1: Bisulfite Conversion***

1. Preheat one thermoshaker to 98°C and set to 1,000 RPM (15s on, 30s off).
2. Preheat another thermoshaker to 54C and set to 1,000 RPM (15s on, 30s off).
3. Set the water bath to 55°C.
4. Thaw urine aliquots and spin down the samples for 5 minutes at 10**°**C and 15,000 RPM.
5. Transfer 500ul of the supernatant to a 15 mL conical tube.
6. Add Lightning Conversion reagent to urine at a 6.5:1 ratio of reagent:supernatant.
7. Incubate in the 98°C thermoshaker for 10 minutes.
8. Incubate in the 54°C thermoshaker for 60 minutes.
9. Immediately proceed to next step

#### ***Step 2: cfDNA Extraction and Desulphonation***

1. Complete the volume to 10 mL using 1x PBS.
2. Add 3 mL Binding Solution K and mix well by vortexing for 10 seconds.
3. Transfer 4.5 mL of the mixture to a MIDI Spin column assembled with one of

the provided collection tubes.

4. Centrifuge for 3 minutes at 1,000 *x g*.
5. Discard the flowthrough and reassemble the spin column.
6. Repeat Steps 3-5 until all the fluid has passed.
7. Apply 3 mL of Wash Solution A to the column and centrifuge for 3 minutes at 1,000 *x g*. Discard flowthrough and repeat.
8. Apply 400 μL of Elution Buffer B to the column let stand at room temperature for 2

minutes. Centrifuge for 2 minutes at 500 *x g*.

9. Reapply the eluted 400 μL Elution Buffer B from Step 8 back to the column and let it stand at room temperature for 2 minutes. Centrifuge for 3 minutes at 500 *x g*.
10. Add 8 μL Proteinase K and mix well by vortexing for 10 seconds, then incubate at 55°C for 10 minutes.
11. After incubation, add 300 μL of Lysis Buffer A and mix well by vortexing for 10 seconds.
12. Add 400 μL of 96-100% ethanol, and mix well by vortexing for 10 seconds.
13. Transfer 750 μL of the mixture from Step 12 into a Mini Spin column assembled with one

of the provided collection tubes. Centrifuge for 2 minutes at 3,300 *x g*.

14. Discard the flowthrough and reassemble the spin column with its collection tube.
15. Repeat Steps 13-14 one more time to transfer the remaining mixture into the Mini Spin column.
16. Apply 600 μL of Wash Solution A to the column and centrifuge for 1 minute at 3,300 *x g*. Discard the flowthrough and reassemble the spin column with its collection tube.
17. Repeat Step 16 one more time, for a total of two washes.
18. Add 200 uL of L-Desulphonation buffer and let it stand at room temperature for 15-20 minutes.
19. Centrifuge for 1 minute at 3,300 *x g*.
20. Add 600 uL of Wash solution A to the column and centrifuge for 1 minute at 3,300 *x g*.
21. Discard the flow-through.
22. Repeat Steps 20-21 for a total of two washes.
23. Spin the column empty for 2 minutes at 14,000 *x g* in a new collection tube.
24. Transfer the column to an elution tube.
25. Apply 30 uL of Elution Buffer B to the column and let stand at room temperature for 2 minutes.
26. Centrifuge for 1 minute at 200 *x g* for 1 minute, and then centrifuge for 2 minutes at 5,200 *x g*.
27. Transfer the eluate back to the column and let stand at room temperature for 2 minutes.
28. Centrifuge for 1 minute at 400 *x g*, followed by centrifugation for two minutes at 5,800 *x g*.
29. Quantify the abundance of the extracted single-stranded cfDNA.

### **Coffee-seq in Plasma**

#### ***Reagents***

- Lightning Conversion Reagent (Zymo Cat# D5032-1)
- L-Desulphonation Buffer (Zymo Cat# D5030-5)
- QIAamp Circulating Nucleic Acid Kit (Qiagen Cat# 55114)
- UDI Primer Set-24 (SRSLY Cat# CBS-UD-24)
- PicoPlus DNA NGS Library Preparation Base Kit (SRSLY Cat# CBS-K250B-24)
- AmpureXP Beads (Beckman Coulter Cat# A63880)
- KAPA HiFi HotStart Uracil+ ReadyMix (2X) (Roche Cat# 7959052001)
- Qubit ssDNA Assay Kit (Invitrogen Cat# Q10212)

#### ***Equipment***

- Two thermoshaker with 15 mL adapters
- Water bath
- Centrifuge

#### ***Step 1: Bisulfite Conversion***

1. Preheat one thermoshaker to 98°C and set to 1,000 RPM (15s on, 30s off).
2. Preheat another thermoshaker to 54°C and set to 1,000 RPM (15s on, 30s off).
3. Set the water bath to 60°C.
4. Thaw plasma aliquots and spin down for 5 mins at 15,0000 RPM and 10°C.
5. Transfer 500 µL of the supernatant to a 15 mL conical tube.
6. Add 600 µL of 1X PBS.
7. Incubate at 98°C for 10 minutes with constant shaking at 1,000 RPM.
8. Centrifuge at 7,500 RPM for 10 minutes.
9. Transfer supernatant to a new 15 mL tube.
10. Add Lightning Conversion reagent to plasma at a 6.5:1 ratio of reagent:supernatant.
11. Incubate in the 98°C thermoshaker for 10 minutes.
12. Incubate in the 54°C thermoshaker for 60 minutes.
13. Immediately proceed to next step

#### ***Step 2: cfDNA Extraction and Desulphonation***

1. Add an appropriate amount of carrier RNA to Buffer ACL.
2. Add 400 µL of Proteinase K into a 50 mL centrifuge tube.
3. Complete the volume to 4 mL with 1X PBS and add to the 50 mL tube containing Proteinase K.
4. Add 3.2 mL Buffer ACL (with carrier RNA). Pulse vortex 30 seconds.
5. Incubate at 60°C for 30 minutes.
6. Place the tube back on the lab bench and unscrew the cap
7. Add 7.2 mL Buffer ACB to the lysate. Pulse vortex 15-30 seconds.
8. Incubate on ice for 5 minutes.
9. Add the mixture to the tube extender and turn on the vacuum.
10. Apply 600 µL ACW1. Drain the column.
11. Apply 750 µL ACW2. Drain the column.
12. Apply 750 µL 96-100% ethanol. Drain the column.
13. Add 200 µL of L-Desulphonation buffer. Close tops and incubate for 15-20 minutes. Drain the column/
14. Apply 750 µL 96-100% ethanol. Drain the column and repeat for a total of two washes.
15. Transfer the column to a collection tube and centrifuge at 14,000 rpm for 3 minutes.
16. Transfer the column to a new collection tube and incubate on a heat block set at 56°C for 10 minutes with the lid open.
17. Place the column in a clean 1.5 mL elution tube. Add 25 µL of Buffer AVE to the center of the column. Close lid and incubate for 3 minutes at room temperature.
18. Centrifuge at 14,000 rpm for 1 minute.
19. Quantify the abundance of the extracted single-stranded cfDNA.

**Sequencing Library Preparation**

Bisulfite conversion of cfDNA involves a cfDNA denaturing step at 98°C such that we get single stranded cfDNA molecules after DNA extraction. For this reason, a single stranded sequencing library preparation method is chosen for the next steps. We prepared sequencing libraries using the SRSLY PicoPlus DNA NGS Library Preparation Base Kit (SRSLY Cat# CBS-K250B-24) with the SRSLY UDI Primer Set-24 (SRSLY Cat# CBS-UD-24) following the manufacturer’s protocol, with the following modifications:

1. The input cfDNA volume used was 18 µL.
2. 1.25 µL of NGS Adapters A and 1.25 µL of NGS Adapters B were added to the 20 µL denatured DNA reaction tube, and the volume was completed by 1.5 µL of ultrapure water.
3. The Index PCR Master Mix was substituted for an equal volume of KAPA HiFi Uracil+ Ready Mix (2X).
4. The Indexed Library DNA Purification step was performed twice, first eluting in 50 µL and then in 25 µL.

### **Bioinformatics Pipeline**

#### ***Sequence data processing and alignment.*** Adapter and low quality bases from the reads were trimmed using BBDuk^1^ (--entropy= ‘0.25’ --maq= ‘10’ -Xmx1g tbo tpe ) and aligned to the C-to-T and G-to-A converted human genome(hg19) using Bismark^2^ with default parameters (Bismark-0.22.1; --unmapped, --quiet). PCR duplicates were removed using Bismark.

***Depth of coverage****.* The depth of sequencing was measured by summing the depth of coverage for each mapped base pair on the human genome after duplicate removal, and dividing by the total length of the human genome (hg19, without unknown bases).

***Removing unconverted molecules.*** Aligned BAM files are filtered to remove unconverted molecules using the Bismark alignment package with default parameters.

***Bisulfite conversion efficiency.*** We estimated bisulfite conversion efficiency by quantifying the rate of C[A/T/C] methylation in human-aligned reads (MethPipe^3^ V3.4.3; -m general -L 500), which are rarely methylated in mammalian genomes.

***Pre-processing of the unmapped reads.*** Reads originating from the Phix genome were removed from the host unmapped reads using Bowtie 2^4^ (--local, --very-sensitive-local, --un-conc). Read IDs from the remaining reads were used to subset paired end reads from the original FASTQ files. Adapter trimming and read quality filtering was performed using BBDuk^1^ (maq=32). Remaining reads were deduplicated using samtools^5^ and merged using FLASH2^6^ (-q -M75 -O). K-mer decontamination to remove human reads was then performed using BBDuk^1^ (k=50

, prealloc = t) and the obtained fastq file was converted to a fasta file for metagenomics analysis.

***Metagenomic abundance estimation from sequencing data.*** Metagenomic analysis of the processed host unmapped reads is performed as previously described^7,8^. Specific to Coffee-seq, read-level filtering of contaminants is performed by removing sequenced reads with more than four cytosines present, or one methylated CpG dinucleotide (the latter represents unmapped, human-derived molecules). Species-level filtering based on the distribution of mapped reads is carried out by first aligning filtered and unfiltered datasets independently. Cytosine-densities of mapping-coordinates in both datasets are measured using custom scripts, and their distributions are compared using a Kolmogorov-Smirnov test. Significantly different filtered-unfiltered distributions are further processed (D-statistic > 0.1 and p-value < 0.05). Briefly, filtered datasets whose distribution of cytosines at mapped locations is significantly lower than unfiltered datasets have one read removed, and are re-tested for differences in their distribution. If the distributions are more similar (as measured through the same criteria), it is filtered out. This process is repeated until one of three criteria is reached: 1) distributions are no longer significantly different, 2) if all reads are removed, or 3) if the D-statistic is small enough. Metagenomic abundances of filtered datasets are estimated using GRAMMy as previously described in Ref 7,8**.** Microbial abundance in downstream analyses was quantified as Molecules Per Million reads (MPM):

*MPM*$= \frac{Adjusted Blast hits x {10}^{6}}{Total Trimmed Reads}$

#### ***Identification of translocated gut bacteria in plasma.*** Fecal shotgun metagenomic data was trimmed using Trim Galore (--nextera --paired) and aligned to the human genome using Bowtie 2^4^ (Bowtie 2.4.3; --maxins 700 --no-discordant --score-min L,0,-0.2). Reads that did not align to the human genome were extracted and assembled using SPAdes^9^ (SPAdes 3.15.3; --meta). The assembled metagenomes were classified using Kaiju^10^ (Kaiju 1.7.4). Paired cfDNA reads obtained after Coffee-seq filtering were then aligned to the assembled metagenomes using Bismark (Bismark 0.22.1).
